# HBV seroepidemiology data for Africa provides insights into transmission and prevention

**DOI:** 10.1101/654061

**Authors:** Anna L McNaughton, José Lourenço, Phillip Armand Bester, Jolynne Mokaya, Sheila F Lumley, Donall Forde, Tongai G Maponga, Kenneth R Katumba, Dominique Goedhals, Sunetra Gupta, Janet Seeley, Robert Newton, Ponsiano Ocama, Philippa C Matthews

## Abstract

**I**nternational goals for elimination of hepatitis B virus (HBV) infection set ambitious targets for 2030. In many African populations, HBV prevalence remains high (≥8%) despite the roll-out of infant HBV immunisation from the mid-1990’s onwards. Enhanced efforts are now urgently required to improve an understanding of population epidemiology, in order to determine which interventions are most likely to be effective in advancing populations towards elimination goals. In populations with a high prevalence of infection, catch-up HBV vaccination of adults has sometimes been deployed as a preventive strategy. An alternative approach of ‘test and treat’ could be applied as a tool to interrupt transmission. We used a systematic approach to investigate the relationship between prevalence of HBV infection (HBsAg) and exposure (anti-HBc) in Africa, and then applied a mathematical model to investigate the impact of catch-up vaccination and a ‘test and treat’ strategy in Uganda, representing a high prevalence setting. We demonstrate a strong relationship between the prevalence of HBsAg and anti-HBc (p<0.0001), but with region-specific differences that may reflect different patterns of transmission. In high prevalence settings, catch-up vaccination may have a transient effect but this intervention does not contribute to a sustained decline in prevalence. In contrast, diagnosing and treating infection has a marked impact on reducing prevalence, equivalent to that of infant immunisation. Conclusion: We have developed a high-resolution picture of HBV epidemiology across Africa. Developing insights into regional differences provides an evidence base for the most effective interventions. In combination with robust neonatal immunisation programmes, testing and treating infection is likely to be of most impact in making advances towards elimination targets.

## INTRODUCTION

There is an estimated global burden of 290 million cases of chronic hepatitis B virus (HBV) infection (1), the majority of which are undiagnosed and untreated (2). Prevalence of HBV exposure and infection can be extremely high in some settings in Africa. For example, in regions of South Sudan and Northern Uganda, seroprevalence of hepatitis B surface antigen (HBsAg) is estimated at 20-25% (3, 4). High endemicity in such settings can be difficult to tackle, as infection can persist for decades, and a persistent population reservoir includes individuals with high viral loads (often corresponding to those with positive HBV e-antigen (HBeAg) status). Furthermore, robust epidemiology data are lacking, and populations in Africa may have vulnerabilities associated with poverty, stigma, and co-endemic human immunodeficiency virus (HIV) infection (2). Horizontal transmission within households, particularly affecting young children, is reported as a significant acquisition route (5-7), but the specific routes and timing of transmission remain uncertain for many African populations.

Vaccination to protect against HBV infection is a cornerstone of interventions aiming to curtail this major public health threat, with enhanced efforts arising as a result of United Nations Sustainable Development Goals (SDGs) setting out elimination targets for the year 2030 (8). HBV vaccination is included in the Expanded Programme for Immunization (EPI), and has been progressively rolled out for infants across southern Africa since 1995. Interventions to prevent mother to child transmission (PMTCT) of HBV infection include accelerated infant vaccination (including a birth dose), combined with antiviral treatment of high risk mothers, and HBV immune globulin (HBIg), if available (9). Routine infant vaccination and enhanced PMTCT regimens currently offer the most likely route to population elimination. However, despite two decades of vaccine implementation, HBV remains endemic in many regions, with time-scales for success that are substantially beyond the SDG targets for 2030 (10, 11).

Tackling the large population reservoir of infection in adults is important, and various strategies are employed to reduce new incident infections in adults, working alongside the established vaccination and PMTCT interventions aimed at children. Introducing ‘catch-up’ vaccination campaigns in older children and adults can appear an attractive public health response in high prevalence settings (12), and this has been undertaken in some settings, despite evidence to suggest a limited population benefit (10) and lack of endorsement by routine recommendations (9). Economic analyses have reported that catch-up HBV vaccine campaigns in young adults are cost-effective only if combined with screening (13), highlighting the importance of focusing not only on prevention but also on investment in diagnosis and treatment (14). The latter concept has been embraced for HIV under the banners of ‘Treatment for Prevention’ (originally coined ‘T4P’) and more recently ‘Universal Test and Treat’ (15), where antiretroviral treatment (ART) is recognised for its role in conferring benefits both to the individual being treated, and also to population health by reducing the risk of transmission (16-18).

Building on this experience from HIV, a ‘test and treat’ HBV strategy could offer substantial advantages. The feasibility, acceptability, and public health consequences of this approach have been positively evaluated through studies in The Gambia (14, 19). Optimisation of vaccine deployment, and evidence-based consideration of the impact of parallel interventions, are urgently required if we are to accelerate progress towards elimination targets in neglected, high-prevalence, resource-limited settings. Here, we used a systematic approach to investigate the sero-epidemiology of HBV across the African subcontinent, based on the principle that understanding the distribution of both active HBV infection and exposure to infection could be of substantial influence in highlighting regional differences and informing the best choice of interventions, especially in situations where resources are limited. Recognising that catch-up vaccine campaigns are being deployed in some locations, we considered the evidence for any benefit, and also assessed the potential impact of a ‘test and treat’ approach. We used an existing model to project the influence of each of these public health interventions in high prevalence settings. Our results have immediate potential for clinical and public health practice, aiming to inform the optimum deployment of limited resources for HBV diagnosis, treatment and prevention.

## METHODS

### HBV seroepidemiology for Africa

We set out to determine the relationship between the prevalence of active HBV infection (HBsAg) and the prevalence of exposure to infection (anti-HBc), through a systematic review of serological data from the published literature. We undertook a systematic search of PubMed and Web of Science in June 2018, using PRISMA criteria (Suppl Fig 1). We used the search terms “HBV antibody”, “anti-HBc”, “HB core antibody”, “HBV exposure” or “HBV prevalence” AND “Africa” or [Name of specific country], using the list of countries on the United Nations (UN) geoscheme for Africa (https://unstats.un.org/unsd/methodology/m49/).

**Fig 1:**
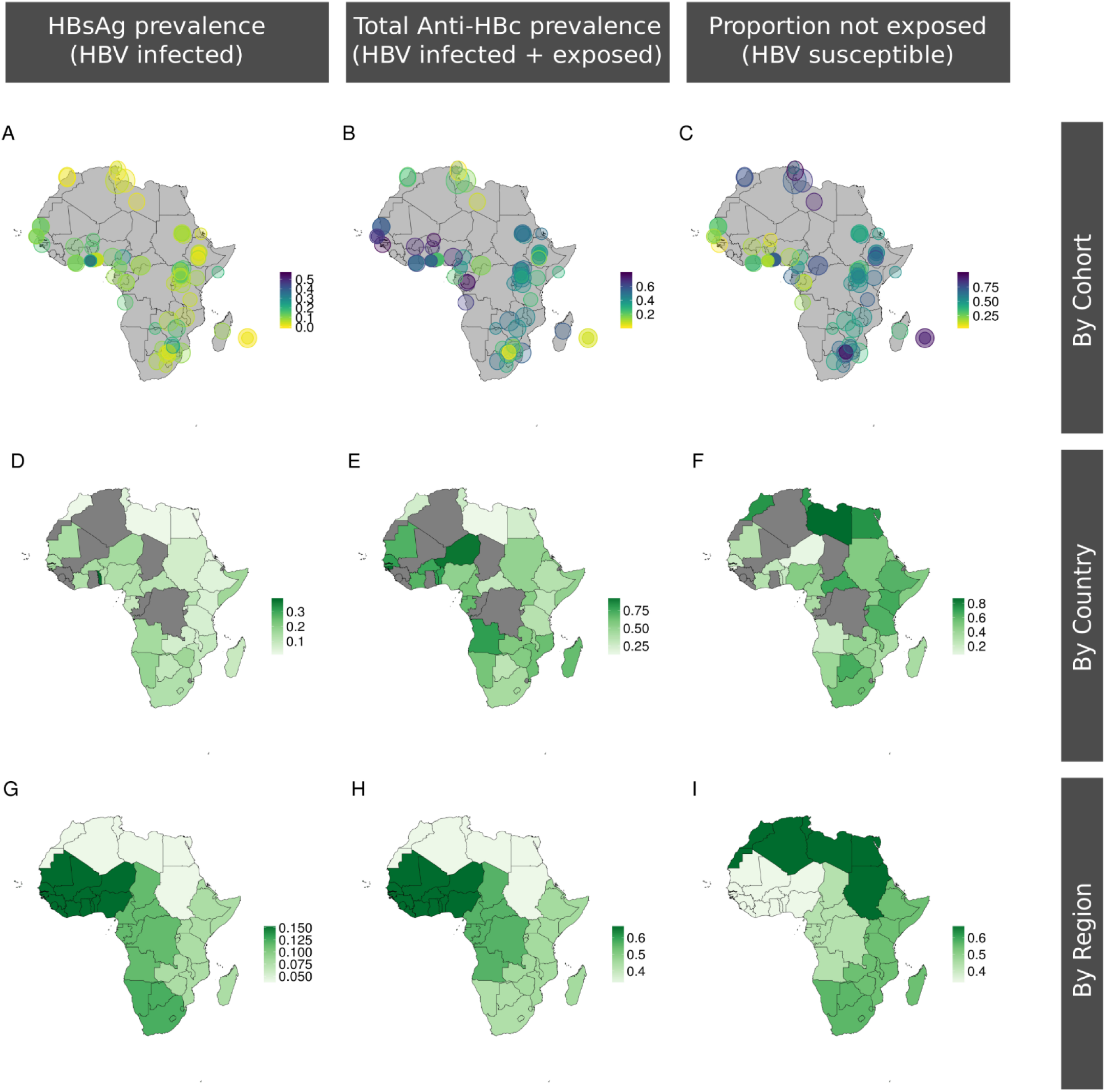
Maps demonstrating the location and HBV seroepidemiology of adult cohorts identified through a systematic literature review. First row shows data by individual cohort, depicting (A) HBsAg prevalence, (B) total anti-HBc prevalence, and (C) HBV susceptible population (100% of population minus anti-HBc prevalence). Each circle is placed to represent the location of the cohort. Second row shows data by country (D-F), and third row by region (G-I). Each area is coloured to reflect high to low prevalence of the attribute in question (scale bar as shown on each panel). Countries shown in grey have no data. The cohort metadata are available on-line, (21) and an interactive version of these maps can be accessed on line using the following link: https://hbv-geo.shinyapps.io/oxafricahbv/. The source code can be accessed here: https://github.com/ArmandBester/Serology_of_HBV_in_Africa.

Inclusion criteria were as follows:

- Data gathered after the widespread roll-out of infant HBV vaccination in Africa in 1995, in order to provide insights that are relevant in the post-vaccine era;
- No reported data collection undertaken pre-1995;
- Reported prevalence of both HBsAg and anti-HBc among cohorts primarily reporting data for adults (age ≥16 years);
- Cohort does not sample a population enriched for HBV infection (specific exclusions are listed in Suppl Fig 1).

We recorded total anti-HBc prevalence (i.e. proportion of population exposed to HBV, irrespective of chronic infection status, termed ‘total exposure’) and also calculated the proportion of the population with cleared infection (i.e. anti-HBc prevalence minus HBsAg prevalence, termed ‘exposed and cleared’). For studies reporting prevalence data from ≥2 cohorts (e.g. HIV-positive and HIV-negative populations), we recorded these as a single publication but ≥2 distinct data points. Studies in a language other than English were translated using Google Translate (https://translate.google.com/). We considered Uganda as an exemplar setting where HBsAg seroprevalence in adults may reach >20% in Northern regions (3, 20), and where catch-up vaccination has been deployed. We also sought evidence for recommendations underpinning catch-up vaccination of adolescents and adults in Africa cited in PubMed using the search terms ‘hepatitis b virus’ or ‘HBV’, and ‘Africa’ or [individual country name], with ‘vaccin*’ and ‘catch up’ or ‘adult’.

Ethics approval was not required for this study, as we analysed data that are already available in the public domain.

### Statistical analysis of metadata

The UN geoscheme classifies Africa into Central, Eastern, Northern, Southern and Western regions; this is a standard approach for sub-dividing macro-geographical areas for statistical analysis. For the regional analysis, each study was assigned equal weighting when analyzing the data, regardless of the study size. We analysed prevalence data for anti-HBc and HBsAg using Graphpad Prism v7.0. For non-parametric data, we sought significant differences between data sets using Mann-Whitney U tests, and for multiple comparisons we used 1-way ANOVA. We used linear regression to derive lines of best fit, 95% confidence intervals and to interpolate HBsAg prevalence from anti-HBc prevalence. We generated maps to illustrate the location of the HBV cohorts and seroprevalence of relevant markers using R (Source code will be made available on acceptance at the following link: https://github.com/ArmandBester/Serology_of_HBV_in_Africa).

### Modelling the impact of adult vaccination vs. ‘test and treat’

In this study we adapted a published dynamic model and Bayesian Markov Chain Monte Carlo approach that we previously developed to fit the seroepidemiology of a population in South Africa, projecting the impact of interventions in that transmission setting (10). As these methods are already published, we have not replicated them in this paper. For ease of reference, we have provided a summary overview of the model population classes and parameters in Suppl Table 1. In this instance, we fitted the model to data from Uganda (Suppl Table 1), in order to represent a setting of high HBsAg prevalence (3).

**Table 1:** Factors that may contribute to regional differences in prevalence of anti-HBc and HBsAg across Africa.

| **Factor** | **Rationale for contribution to regional differences in HBV seroepidemiology** |
| --- | --- |
| **Circulating HBV viral genotype** | Predominant genotype varies by region with genotype-A common in Southern Africa, genotype-D in the North and genotype-E in the West (39). |
| **Host ethnicity and genetics** | HLA-type and T-cell repertoire have been linked to the ability to control the infection (40-42). |
| **Transmission differences** | Subtle differences in the transmission patterns (vertical vs horizontal) of the HBV genotypes have been documented. Transmission route is fundamentally linked to age at exposure (43). |
| **Age at exposure** | The probability of developing chronic HBV after exposure is strongly associated with age (44). Populations with a younger age at exposure are therefore likely to have a higher HBsAg prevalence relative to the anti-HBc prevalence (Fig 5A). |
| **Co-infection within population** | Risk factors for acquisition of blood-borne viruses overlap between HIV, HBV and HCV. Egypt and the Nile Delta have some of the highest reported prevalences of HCV globally. Co-infection of HBV and HCV has been linked to spontaneous clearance of HCV although evidence of the impact on HBV remains scarce (45, 46). |
| **Political instability** | Central Africa includes several regions disrupted by recent conflict and resulting population migration, with powerful influence on increases in interpersonal violence and sexual assault, reduced access to barrier contraception, inadequate screening of blood products, and reduced access to healthcare, all of which can increase exposure rates in the adult population. |
| **Traditional cultural practices** | Exposure to blood-borne viruses is influenced by traditional healing practices, scarification, piercing, tattooing and non-sterile surgical practice (e.g. circumcision). |
| **Uptake of HBV vaccination in the region** | Countries with earlier uptake of the HBV vaccine are likely to have lower anti-HBc and HBsAg prevalence than countries that implemented the vaccine later. Prevalence of vaccine escape mutants may contribute, although data for Africa are scarce (33). |
| **Potential role of insect vector** | Biting insects capable of mechanical transmission of HBV may be prevalent in some regions, although there is a lack of firm evidence base for HBV transmission (47). |

We used published seroepidemiology variables, as follows: HBsAg prevalence 10.3%, anti-HBc prevalence 42% and HBeAg-positive (HBeAg+) relative prevalence at 27% (Suppl Table 1), which the model robustly recovered at 10% (95% CI 7.92-11.7%), 42.1% (95% CI 40.2-44.0%), and 26.9% (95% CI 24.8-29.0%), respectively. We left four parameters free to be fitted (vertical transmission rate for HBeAg+ and HBeAg-negative (HBeAg-), rate of conversion from HBeAg+ to HBeAg-, and spontaneous clearance of chronic HBV), for which the posteriors matched literature expectations (Suppl Table 1). PMTCT (combining accelerated neonatal immunisation with HBIg and antiviral therapy in pregnant mothers) and vaccine-based interventions were modelled as previously described (10), and we added a ‘test and treat’ strategy. The latter was simplified to reducing the transmission potential of the HBV infected proportionally to the control effort (e.g. 20% coverage of test and treat in a particular age-group equated to a 20% reduction in that group’s force of infection).

## RESULTS

### Significant relationship between prevalence of HBsAg (infection) and anti-HBc (exposure)

Through a systematic literature review, we collated prevalence data for HBsAg and total anti-HBc, identifying a total of 88 studies spanning 37 African countries and generating 100 unique data points (complete metadata are available on-line) (21). Information on studies reporting prevalence data from ≥2 cohorts (n=12) is recorded in Suppl Table 2. The median ages for the cohorts represented was 34.4 years (IQR 29.1-36.2 years) based on age data available for 64% of studies.

The distribution of these cohorts and the prevalence of HBV serological markers is shown in Fig 1. These data can be interactively explored on-line at https://hbv-geo.shinyapps.io/oxafricahbv/. Pooling data for all regions, the prevalence of HBsAg (infection) was positively correlated with total anti-HBc (exposure), R^2^=0.35, p<0.0001 by linear regression, Fig 2A. Median HBsAg prevalence across Africa was 9.3% (IQR 5.5-15.1%) with an anti-HBc prevalence of 53.0% (IQR 34.4-69.2%). We did not find any significant differences in HBsAg or anti-HBc prevalence between HIV+ cohorts (N=26) and all other cohorts (N=74; p=0.16 and p=0.42, respectively; Suppl. Fig 2).

**Fig 2:**
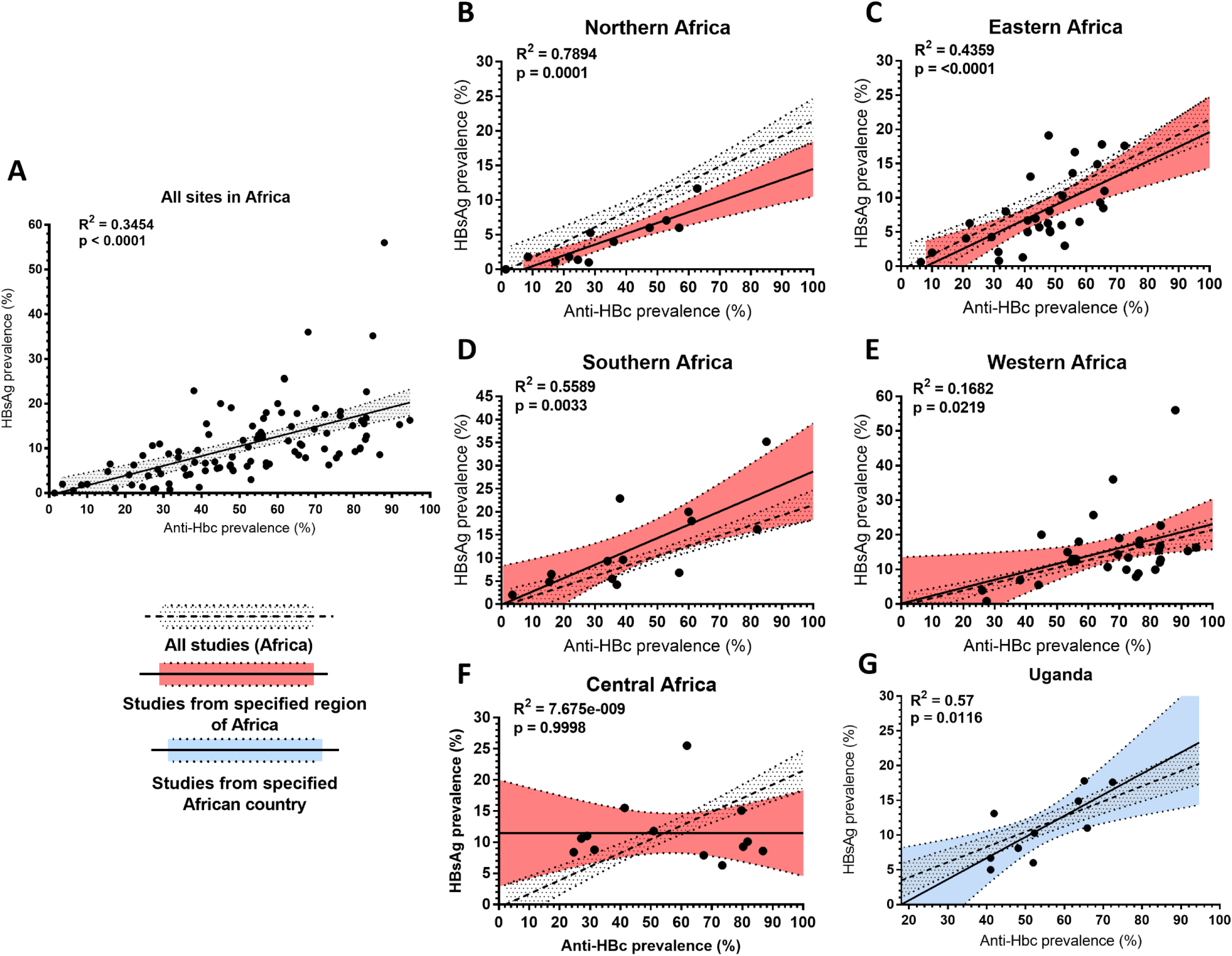
Relationship between population prevalence of anti-HBc (exposure) and HBsAg (active infection) for different regions of Africa. Data are shown for (A) the entire African sub-continent, (B) Northern (C) Eastern (D) Southern (E) Western (F) Central, (G) Uganda. These data are derived from a review of the published literature (full metadata available on-line)(21). The UN geoscheme used to classify the geographic regions can be found at https://unstats.un.org/unsd/methodology/m49/. R^2^ and p values by linear regression (solid line). Outer dashed lines show 95% confidence intervals. Linear regression plots and 95% confidence intervals (shaded regions) are shown for the whole of Africa in grey, for each region in red, and for a single country in blue. Data in plots B-G have been shown together with data for the whole continent for comparison.

### Variations by region and by country

For most regions, we observed the same overall association between total anti-HBc and HBsAg, Fig 2B-E, but with some interesting variations. Northern Africa has lower prevalence rates of infection than other regions (Fig 2B and Fig 3A,B). In contrast, Western Africa has the highest population exposure and correspondingly highest rates of HBsAg positivity (Fig 2E, Fig 3). HBsAg prevalence differs significantly between regions (for Northern Africa compared to Western and Southern Africa, p=0.0002 and p=0.04 respectively, Fig 3B); and cannot be explained only by lower population exposure rates: although anti-HBc prevalence is somewhat lower in Northern than Western Africa (p=0.001), there is no difference in anti-HBc prevalence between Southern and Northern Africa (p=0.99). Indeed, the predicted HBsAg prevalence was approximately 50% lower in Northern than Southern Africa for any given anti-HBc prevalence (Fig 2B, 2D; Suppl Table 3).

**Fig 3:**
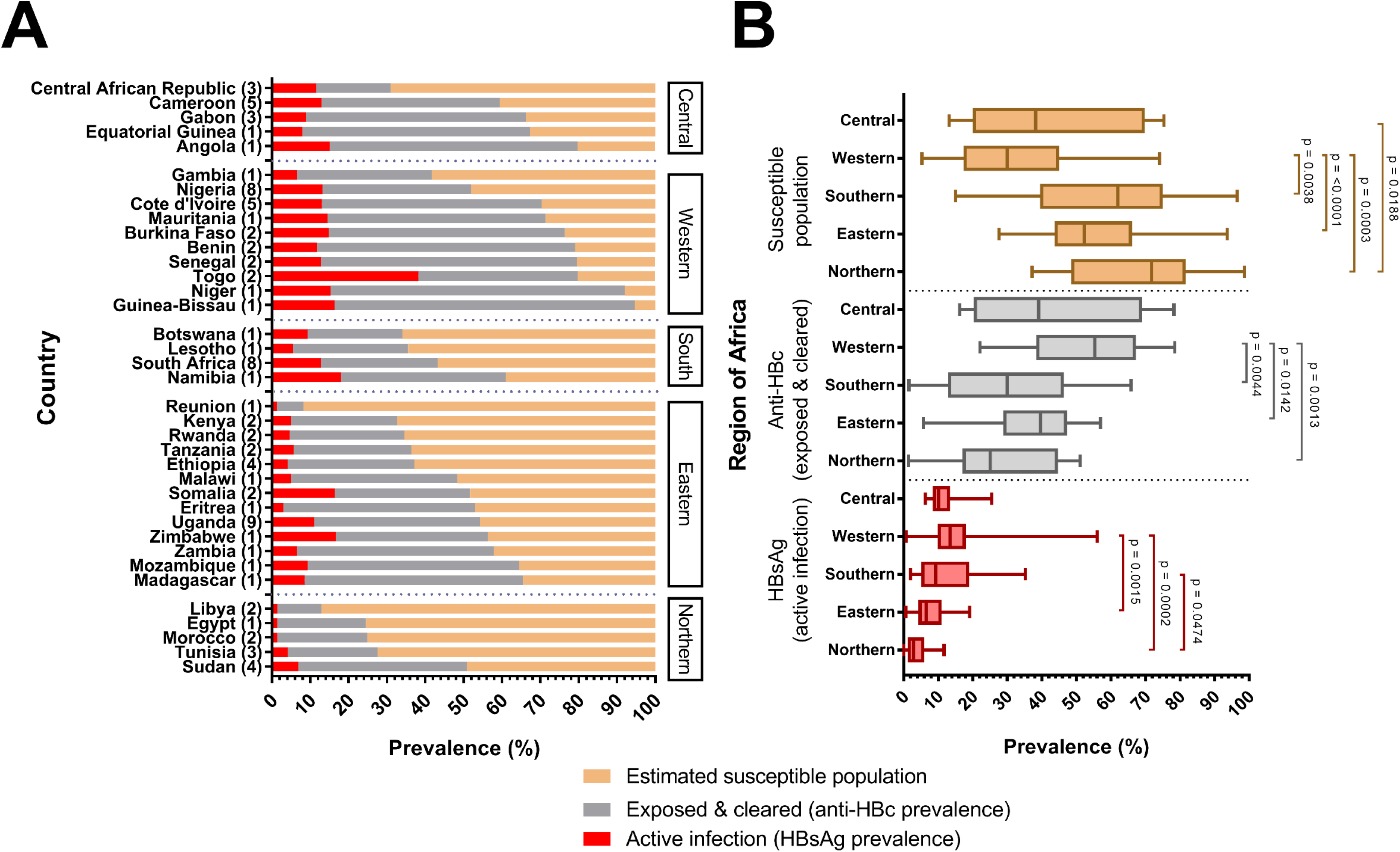
Estimated proportion of the population with active HBV infection, previous exposure and susceptibility to HBV infection, divided by (A) country and (B) region of Africa. Countries have been grouped by region according to the UN geoscheme for Africa. The number of studies per country is given in brackets next to the country name. Two studies were counted twice as they contained cohorts from two different countries. In (B), boxplots show the mean, inter-quartile ranges and range of the data sets, with all significant differences indicated. All studies are listed in on-line metadata.(21) See methods for definitions of infection, previous exposure and susceptibility.

Central African regions display a different relationship, whereby high population HBV exposure is not associated with a correspondingly high prevalence of infection (Fig 2F). This is likely to be a robust representation of the region, as the data cover a 15-year period, and represent multiple countries from where a median of 455 subjects were analysed (IQR 225-782 subjects). Focusing specifically on Uganda, in Eastern Africa, we also found a significant relationship between HBsAg and anti-HBc prevalence; p=0.01, Fig 2G, 3A. However, even within this single country, considerable differences are seen in the ratio of HBsAg:anti-HBc between different studies (see metadata on-line (21)).

In three studies that assessed both HIV-positive and HIV-negative cohorts, HBsAg prevalence was higher among HIV-positive subjects (mean 2.23-fold) (22-24). Anti-HBc prevalence was also higher in HIV-positive cohorts than in HIV-negative cohorts for 2/3 studies (22, 24). In a third study of highly exposed cohorts in South Africa, anti-HBc prevalence was similar irrespective of HIV status, suggesting the increased HBsAg prevalence in the HIV-positive cohort was the result of reduced clearance rates relative to the HIV-negative cohort (23).

### Impact of catch-up vaccination of adolescents and adults

We did not identify any published evidence or specific recommendations for catch-up vaccination of adolescents and adults, either in the form of intervention studies or literature reviews. However, a number of authors do suggest catch-up vaccine programmes as a way of tackling high population HBV prevalence (3, 12, 25, 26) (data from literature review summarised in Suppl Table 4).

Based on combining the mean prevalence values from Uganda cohorts to provide a broad overview, 54% of adults across this country have been exposed (among these, a total of 11% of adults are actively HBV-infected, and the remainder have been infected and cleared). This leaves 46% of the total adult population potentially susceptible (orange bars, Fig 3A). Only a small proportion of this susceptible pool of adults would be exposed to infection each year (there are few data to estimate this exposure rate, but one study from another region of East Africa estimates this at 3-4%) (27). The natural history of HBV infection in adults suggests that <5% of exposure events lead on to chronic infection. Thus, the predicted proportion of the total adult population predicted to avoid chronic infection through catch-up vaccination each year is, roughly, 50% (vulnerable) × 4% (exposed) × 5% (develop chronicity) = 0.1%.

Using our established model of HBV transmission and prevention (10), we investigated the impact of catch-up vaccination among adults within a high HBV prevalence setting, exemplified by Uganda (3) (Suppl Table 1). Selected results from simulations are presented in Fig 4, in which a catch-up immunization programme in adults is projected to have only a transient impact on reducing new cases of HBV infection. In the long-term, this strategy offers no sustained overall benefit in progress towards elimination targets, even when deployed at 100% population coverage (Fig 4A, orange band). The poor impact of catch-up vaccination, estimated at only an 8% reduction over 200 years (Fig 4A), is due to a limited pool of susceptible adults and the lack of impact on the actively infected population. In contrast, enhanced coverage of other interventions, including PMTCT and infant immunisation will lead to shorter time-frames for reducing HBsAg prevalence, given their direct impact on the rate of new chronic infections, the main reservoir of HBV infection.

**Fig 4:**
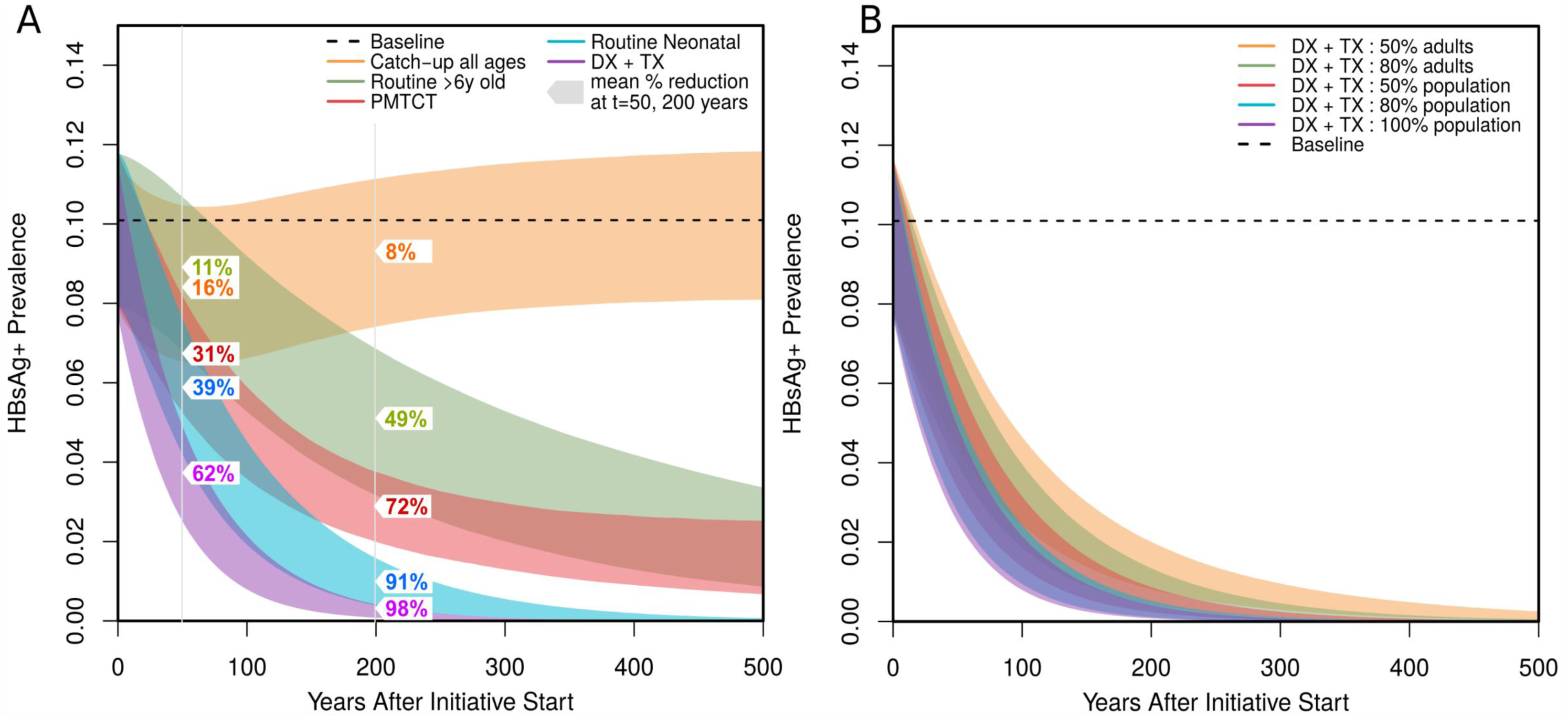
Simulation of change in HBsAg prevalence over time in response to population interventions. The pre-intervention prevalence is set close to 10%, based on population prevalence of HBV infection in Uganda (Suppl Table 1). Decline in prevalence is shown over time; bands are 95% CI for each intervention based on 5000 stochastic simulations using parameter samples from the posteriors obtained by fitting the model. (A) Comparison of interventions applied to 100% of the population: catch-up vaccination of all ages as a one-off event at time=0 (orange), routine immunisation of children aged >6 years as an alternative catch-up strategy (green), PMTCT all births (combining accelerated neonatal immunisation with HBIg and antiviral therapy in pregnant mothers, red), routine neonatal immunisation (blue) and diagnosis and treatment ‘Dx + Tx’ (purple); (B) Comparison of Dx + Tx applied to different proportions of the population: 50% of adults (orange), 80% of adults (green); 50% of whole population (red); 80% of whole population (blue); 100% of whole population (purple). Fitted baseline prevalence is indicated by the dashed line. All interventions modelled as previously described (10), and the new ‘Dx + Tx’ is simplified to a reducing the force of infection of each population group by the specified amount (see main text).The numbers at time points t=50 and t=200 years are the mean reduction in HBsAg prevalence achieved for each of the interventions.

### Impact of ‘test and treat’ in highly endemic settings

We also modelled the impact of ‘test and treat’, based on the premise that the whole population is screened, projecting that this strategy has the fastest reduction in HBV population prevalence of all interventions with 62% reduction in prevalence by 50 years, and 98% at 200 years (Fig 4A, purple band). Recognising the significant barriers to identifying all cases of infection, (including silent infection, lack of education, poor access to laboratory facilities, and stigma) (2), we also modelled the outcome for ‘test and treat’ strategies that reach <100% of the HBV-infected population. Diagnosis and treatment for 80% of infected adults (Fig 4B, green band) or 50% of the whole infected population (Fig 4B, red band) delivers a reduction in HBsAg prevalence over time that is comparable to infant immunisation (Fig 4A, blue band). Even reducing the population tested and treated to only 50% of adults (Fig 4B, orange band) is still substantially more effective than 100% catch-up vaccination (Fig 4A, orange band).

## DISCUSSION

United Nations Sustainable Development goals have set an ambitious time-frame in which to make significant reductions in both prevalence and incidence of HbsAg carriage by the year 2030 (8). Careful, evidence-based deployment of interventions is essential if sustained and collective progress is to be made towards these targets. We have here shown how existing epidemiology data can provide important insights into patterns of infection and susceptibility. Other systematic reviews and global estimates of HBsAg prevalence have been published over the last few years (1, 4, 28); our approach differs in also accounting for the prevalence of exposure, and in considering the relationship between infection and exposure in different settings.

Although it can seem intuitive to deploy catch-up vaccination for adolescents and adults in high prevalence HBV settings, we here demonstrate that only a limited proportion of individuals remain susceptible in these populations, representing a minority who will potentially benefit from catch-up vaccination. The effectiveness of catch-up vaccination is strongly linked to the size of the susceptible population, as illustrated by Fig 5B, with a greater impact seen in low-prevalence populations. For this reason, catch-up vaccination will frequently not be a prudent use of resources, although in some settings, there may be cost benefits in targeting young populations with catch-up vaccination (29). The distinct regional patterns of HBV epidemiology, and the lack of overlap between the epicentres of HCV infection in North Africa, HIV in Southern Africa and endemic HBV, suggest different patterns of transmission of HBV between regions, and different transmission routes for different blood-borne viruses across the continent. Notably, even with a single country – exemplified here by Uganda – there is evidence of region-specific differences in exposure and transmission.

**Fig 5:**
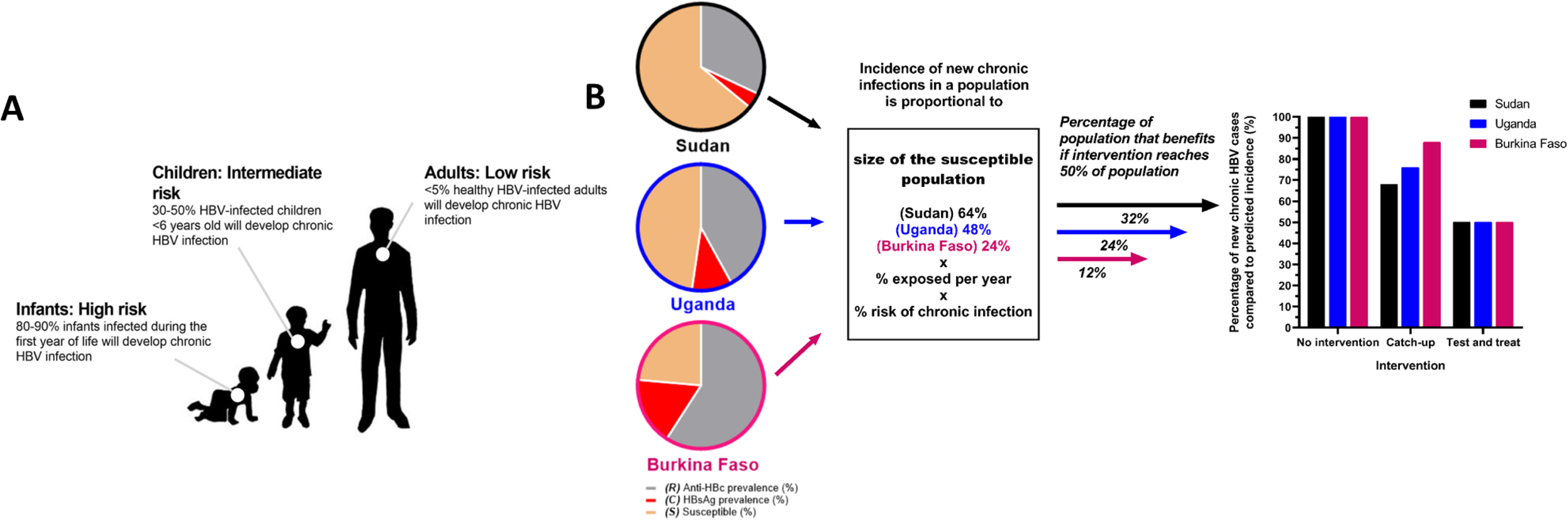
Cartoons to illustrate seroepidemiology of HBV infection in Africa and the differential impact of HBV interventions according to population targeted. (A) After exposure to HBV, the risk of developing chronic infection is highest amongst young infants and this risk gradually declines with age until adulthood, where there is low risk of developing of chronic infection. Figure informed by parameters in Suppl Table 1. (B) Populations from Sudan (48) Uganda (3) and Burkina Faso (49) represent the 25th, 50th and 75th percentiles in the data set collected from our literature review. In adults, assuming that different populations are exposed at the same rate and the risk of chronic infection is constant (estimated to be 5% in healthy adults as shown in Fig 5A), the incidence of new chronic HBV infection in the population is related to the susceptible proportion (*S*). Without intervention, 100% of predicted new cases will occur. If 50% of the adult population is vaccinated in a catch-up campaign, chronic infection will be prevented only among the population *S*. The impact of catch-up vaccination on incidence is therefore related to *S*, with reduced impact in highly exposed populations. In a test and treat scenario, with 50% of cases identified and treated, incidence is consistently reduced, regardless of *S*.

In order to make progress towards HBV elimination goals, we therefore suggest that the public health agenda should prioritise active ‘test and treat’ programmes aimed at older children and adults. Success of this strategy depends on education, resource and infrastructure. Our results are congruent with the findings of a recent review of HBV vaccination in South Africa highlighting the need to prioritise infant immunization above catch-up campaigns in adolescents (26), and with previous economic evaluations of the ‘test and treat’ approach (30, 31). In practice, achieving success through ‘test and treat’ requires multi-pronged investment including education, laboratory infrastructure to provide assessment and monitoring of infection, and provision of effective, sustained drug therapy for both HBV monoinfection and HIV/HBV coinfection. In order for treatment to be successfully rolled out, focus on diagnosis is pre-requisite (14, 32), parallel investment in infra-structure is paramount to triage cases for treatment (based on including laboratory and radiological criteria), and additional scrutiny will be required for drug resistance (33).

The epidemiology and dynamics of infection are different in certain high-risk subgroups (health care workers, partners and household contacts of infected individuals, sex workers and their clients, men who have sex with men), and continuing to target these individuals with preventive vaccination remains very important. Likewise, we continue to emphasise the importance of routine infant immunization campaigns which are a cornerstone of elimination strategies (10).

### Relationship between exposure and active HBV infection in Africa

Our seroepidemiology review highlighted considerable regional differences in the relationship between HBV exposure and active infection. A diverse range of factors influence the risk of developing chronic HBV infection after acute infection (Table 1), with age at exposure among the most robustly recognised. Our data suggest that in regions with low HBsAg prevalence in the setting of high anti-HBc (epitomised by countries in central Africa), most exposure events may be occurring in adults. In contrast, in Western Africa, where HBsAg prevalence is highest, the majority of exposure events may be in early life. Careful data collection and review is required to underpin the most effective interventions for specific locations.

Genotype of infection and transmission routes should also be considered as factors influencing sero-epidemiology. HBV genotypes A, D and E are most prevalent in Africa, with a substantial proportion of infections accounted for by horizontal transmission during early childhood (34). Data remain scarce but, an increased HBV HBeAg prevalence amongst genotype E infected individuals has been reported (35), typically correlating with higher viral loads and increased risk of vertical transmission (36). Genotype E is geographically restricted to Western Africa, where we describe the highest HBsAg prevalence, suggesting infection in this region may be occurring at an earlier age than elsewhere. Likewise, traditional cultural practices that confer exposure to HBV at specific ages may be common in some regions but not others. Scarification has been correlated with increased HBV risk in Nigeria (37), and unsafe medical practices and a lack of awareness of risk factors for HBV may contribute towards transmission in some populations.

### Relationship between HBV and HIV

There was no evidence from our dataset that HIV+ individuals were more likely to be either HBV infected or exposed, in keeping with previous reports (36). This observation reflects different transmission patterns: HIV is less infectious than HBV when transmitted by blood and is largely sexually transmitted in Africa. In contrast, the risk of developing chronic HBV infection is high in early life and declines with age. However, robust analysis of the influence of HIV on HBV exposure and acquisition is made difficult by limited data. While we were able to identify a large number of HIV+ cohorts, only three of these had directly comparable HIV-negative cohorts (data from South Africa and Uganda) (21). Among all other published cohorts, which we have assumed to be HIV-negative, a background prevalence of HIV infection is likely but not clearly reported.

### Caveats and limitations

Given Africa’s population of >1.2 billion people and the substantial public health problem that HBV represents for this continent, there are very limited epidemiological data to inform the most appropriate interventions. Our maps highlight geographical gaps in the data (Fig 1), while existing cohorts are often relatively small and biased by the recruitment of specific groups who may not be representative of the general population. The published literature does not account for the prevalence of occult HBV, which is rarely detected due to lack of availability and high cost of HBV DNA testing. However, individuals with occult HBV would still generate anti-HBc; thus while we may be underestimating the prevalence of active infection, these subjects are still included within our exposed population.

We did not include data for anti-HBs prevalence (immunised population) as a limited number of papers report the prevalence of anti-HBs together with anti-HBc and HBsAg data. The most common reason for study exclusion form the literature review was no anti-HBc prevalence reported (Suppl Fig 1). Making the inclusion criteria more stringent would have limited the findings from the study.

We included papers published after the EPI introduction of HBV vaccine in 1995, in order to make our study applicable to current-day vaccinated populations, although in practice, roll-out of the vaccine was patchy and adopted at a variable rate over the decade that followed. There are limited data for many regions describing the prevalence of three-dose vaccine coverage. Based on the age of adults represented in most of our cohorts, we can assume the majority of subjects in the study were unlikely to have been vaccinated at birth. Future sero-surveys will provide more insights into the impact of routine infant HBV vaccination. An assessment of vaccine-mediated immunity (anti-HBs) would also be useful in estimating the impact of infant HBV vaccination in Africa.

For this study, we focused on adult populations only, as the age-associated risk of developing chronic HBV is a confounding factor in younger cohorts, making inference about the anti-HBc prevalence challenging across multiple age groups. It would be of interest to determine age-specific prevalence of HBsAg and anti-HBc because age is likely to be an important source of heterogeneity. However, metadata are poorly reported by existing literature and we were unable to disaggregate serological data by age.

Our dynamic model includes a series of simplifying assumptions. For instance, our ‘test and treat’ intervention does not stratify individuals for therapy, but works on the basis of treating anyone who is HBsAg-positive. In current clinical practice, guidelines recommend treatment in the context of high viral load and/or evidence of inflammatory liver disease (38). However, explicitly stratifying population subgroups for ‘test and treat’ within our framework would have required the inclusion of epidemiological classes (e.g. clinical progression or population classes stratified by viral load or liver transaminases), which would have added significant uncertainty to our projections. Our model framework does not include explicit age-specific or risk-group assumptions regarding force of transmission, and again we argue that little data exists to inform this parameterization and adds extra classes with added uncertainty. Keeping parameterisation simple was an intended approach, as is general practice in dynamic modelling. Our projections are not intended to be exact quantifications of impact over time, but serve as means of comparing the dynamic and non-linear outcomes of different strategies.

### Implications for practice

An improved understanding of HBV epidemiology at local and regional levels will be informative for the design of public health initiatives, allowing relevant, targeted interventions to be deployed in individual settings. Catch-up vaccination is not routinely endorsed by guidelines, but is nevertheless being deployed by some public health initiatives devised in response to high prevalence settings. Our data show the added value of ‘test and treat’ approaches for HBV, building on experience gained from HIV. We advocate significant investment in capacity building for improving HBV diagnosis and treatment, including point-of-care testing, antenatal screening, and provision of TDF. A sustained and systematic commitment to diagnosis and treatment represents a key component of the journey towards HBV elimination.

## Abbreviations

Anti-HBc: antibody to hepatitis B core antigen
ART: antiretroviral therapy
EPI: Expanded Programme for Immunization
HBeAg: Hepatitis B e-antigen
HBIg: Hepatitis B immunoglobulin
HBsAg: Hepatitis B surface antigen
HBV: Hepatitis B virus
HIV: Human immunodeficiency virus
IQR: Interquartile range
PMTCT: prevention of mother to child transmission
SDGs: Sustainable Development Goals
UN: United Nations

## SUPPLEMENTARY DATA

Full metadata for our systematic literature review are available on-line (21).

**Suppl. Table 1: Population data and HBV seroepidemiology for Uganda used to inform a model to determine impact of interventions.** Further details of the model have been previously described (10).

**Suppl. Table 2: Details of studies from Africa reporting HBV prevalence data from ≥2 cohorts.** These studies (n=12) were recorded as a single study but ≥2 data points (as appropriate). Differences in the cohorts are highlighted in city/location, cohort characteristics and cohort size. Complete metadata for the manuscript are available at https://figshare.com/s/4414fce1d474bc8a6198.

**Suppl. Table 3: Predicted HBsAg prevalence for Northern, Eastern, Southern, Western and Central Africa with a given anti-HBc prevalence.** Data to inform the analysis were derived from a systematic literature review (full metadata on-line)(21). Linear regression analysis data for the African regions was simulated to predict HBsAg prevalence with a given anti-HBc prevalence ranging from 5-60% and increasing in increments of 5%. Values plotted in Suppl. Fig 3.

**Suppl. Table 4: Results of a systematic literature review to identify evidence or recommendations for use of catch up HBV vaccination in adolescents and adults in Africa.**

**Suppl. Fig 1: PRISMA chart to show the search criteria and relevant literature identified through a systematic literature review to describe the relationship between the prevalence of HBsAg and anti-HBc in subSaharan Africa**. The resulting metadata set is available on-line (21).

**Suppl. Fig 2: Average prevalence of anti-HBc and HbsAg in confirmed HIV-positive cohorts and all other cohorts based on data for Africa collected through a systematic literature review.** Boxplots show the mean, inter-quartile ranges and range of the data sets. No significant differences were identified for either anti-HBc or HBsAg prevalence (p=0·42 and 0·16 respectively).

**Suppl. Fig 3: Predicted HBsAg prevalence for Northern, Eastern, Southern, Western and Central regions of Africa with a given total anti-HBc prevalence (reflecting exposure).** Linear regression analysis data for each region was simulated to predict HBsAg prevalence with a given anti-HBc prevalence ranging from 5-60% and increasing in increments of 5%. Plotted from values given in Suppl. Table 2.

## AUTHORS’ CONTRIBUTIONS

The article was conceived and designed by ALM, JS, RN, PO and PCM. The paper and figures were written by ALM, JL and PCM with editorial contributions from all authors. ALM, SFL, JM and DF undertook the systematic literature review. JL and SG provided the mathematical model and simulations, with input from DG. PAB analysed epidemiology data and generated interactive maps. KRK provided expertise in health economics. TGM, KRK, JS, RN and PO provided expertise on local HBV interventions in South Africa and Uganda. All authors approved the final manuscript.

