## Supplementary material for "HBV seroepidemiology data for Africa provides insights into transmission and prevention"

**CONTENTS:**

**Suppl. Table 1:** Population data and HBV seroepidemiology for Uganda used to inform a model to determine impact of interventions: ***page 2-3***

**Suppl. Table 2:** Details of studies from Africa reporting HBV prevalence data from ≥2 cohorts: ***page 4-7***

**Suppl. Table 3:** Predicted HBsAg prevalence for Northern, Eastern, Southern, Western and Central Africa based on a given anti-HBc prevalence: ***page 8***

**Suppl. Table 4:** Results of a systematic literature review to identify evidence or recommendations for use of catch up HBV vaccination in adolescents and adults in Africa: ***page 9-10***

**Suppl. Fig 1:** Flow chart for systematic literature review of studies reporting HBV seroepidemiology data for countries in Africa, based on PRISMA criteria: ***page 11***

**Suppl. Fig 2:** Average prevalence of anti-HBc and HBsAg in confirmed HIV-positive cohorts and all other cohorts: ***page 12***

**Suppl. Fig 3:** Predicted HBsAg prevalence for Northern, Eastern, Southern, Western and Central regions of Africa with a given total anti-HBc prevalence: ***page 13***

**References for suppl material:** ***page 14-17***

**Supplementary Table 1: Population data and HBV seroepidemiology for Uganda used to inform a model to determine impact of interventions.**

Further details of the model have been previously described (1).

| Variable | Model Value (input) | Fitted by Model (output) | Literature Support | Reference | |
|---|---|---|---|---|---|
| HBV+ prevalence (HIV-) | 10.30% | 10.0% (95% CI 7.92-11.7) | 10.30% | Bwogi et al., (2) | HBV prevalence |
| HBV+ prevalence (HIV+) | 10.30% | 10.0% (95% CI 7.92-11.7) | 10.3% | **Presume same as for general population (2). This assumption is previously reported (3) as well as being supported by current literature review. | |
| Anti-HBc prevalence (exposure seroprevalence) | 42.0% | 42.14% (95% CI 40.2-44%) | (Anti-HBc) – (HBsAg+ only) = 52.30% - 10.3%= 42% | Bwogi et al., (2) | |
| HIV prevalence in individuals >6 years of age in Uganda | Fixed at 6.5% (based on adult prevalence) | ----- | 6.5% | UNAIDS data for Uganda, (4) |  |
| HIV prevalence in individuals age 1-6 years in Uganda | Fixed at 0.5% | ----- | 0.7% | Note no estimate in UNAIDS data. Conservative estimate used, based on Uganda AIDS indicator survey (5) |  |
| HIV prevalence in individuals <1 years of age in Uganda | Fixed at 0.5% | ----- | 0.7% | Note no estimate in UNAIDS data. Conservative estimate used, based on Uganda AIDS indicator survey (5) |  |
| Vertical transmission rate for HBV (HBeAg+) | free (uninformative prior) | 80.4% (95% CI 71.9-88.4%) | 70-80% | Gentile & Borgia (6) | HBV transmission |
| Vertical transmission rate for HBV (HBeAg-) | free (uninformative prior) | 24.6% (95% CI 14.7-33.4%) | 10-40% | Gentile & Borgia (6) | |
| HBeAg+ prevalence (within HBsAg+) | 27% | 26.9% (95% CI 24.8-29.0%) | 27% | Matthews et al., (7) | |
| Clearance in individuals >6 years of age (or percent expected to have acute infection) | fixed at 95% | ---- | >95% | WHO HBV factsheet (8) |  |

| | | | | | |
|---|---|---|---|---|---|
| Clearance in individuals <1 years of age (or percent expected to have acute infection) | Fixed at 15% | ---- | 10-20% | WHO HBV factsheet, updated July 2016 (8) |  |
| Clearance in individuals 1-6 years of age (or percent expected to have acute infection) | Fixed at 40% | ---- | 30-50% | WHO HBV factsheet (8) |  |
| Spontaneous clearance of chronic HBV | Free (uninformative prior) | 0.31% (95% CI 0.37-0.71%) | 0.73%; 1.15%; 2.26% annually | Ferreira et al., (9); Chu et al., (10); Lui et al., (11) (respectively) | |
| Duration of an acute infection | Fixed at 6 months | ----- | 6 months | Liang., (12) |  |
| Rate of conversion from HBeAg+ to HBeAg- | Free (uninformative prior) | 5.12% (95% CI 4.25-6.38%) across ages | 0.8% a year for <5 yrs; 3% for 5-15 yrs; 8-15% for >15 yrs | Kao., (13) | |
| Life expectancy | Fixed at 63 years | ------- | Males: 62.2yrs, Females: 64.2yrs | Uganda Bureau of Statistics, (14) | Demographics |
| Efficacy of vaccination against HBV infection if HIV- in ages <1, 1-6, >6 years | fixed at 95.2%, 89.2%, 79.6% | ------ | ---- | Estimated using this model and data from SA in previous study by McNaughton et al., (1) | Vaccine efficacy |
| Efficacy of vaccination against HBV infection if HIV+ in ages <1, 1-6, >6 years | fixed at 78.4%, 21.7%, 0.31% | ------- | ---- | Estimated using this model and data from SA in previous study by McNaughton et al., (1) |  |

**Suppl Table 2: Details of studies from Africa reporting HBV prevalence (prev.) data from ≥2 cohorts**. These studies (n=12) were recorded as a single study but ≥2 data points (as appropriate). Differences in the cohorts are highlighted in city/location, cohort characteristics and cohort size. Complete metadata for the manuscript are available at https://figshare.com/s/4414fce1d474bc8a6198 (15).

| Country | First author and reference | Citation | PMID or DOI | City / location | Cohort Characteristics | Size of cohort (n=) | Anti-Hbc prev. | Anti-HBc (& HBsAg negative) prev. | HBsAg prev. | Estimated susceptible population (100% - anti-Hbc prev.) | Reason for multiple cohorts |
|---|---|---|---|---|---|---|---|---|---|---|---|
| **Burkina Faso** | Collenberg (16) | J Med Virol. 2006 May;78(5):683-92. | DOI: 10.1002/jmv.20593 | Ouagadougou (urban) | Blood donors, antenatal | 238 | 76.40% | 59.10% | 17.30% | 23.60% | Urban (Ouagado-ugou) vs Rural (Nouna) |
| **Burkina Faso** | Collenberg (16) | J Med Virol. 2006 May;78(5):683-92. | DOI: 10.1002/jmv.20593 | Nouna (rural) | Blood donors, antenatal | 289 | 69.60% | 55.30% | 14.30% | 30.40% | |
| **Ethiopia** | Shiferaw (17) | BMC Res Notes. 2011 Nov 3;4:479. | DOI: 10.1186/1756-0500-4-479 | Addis Ababa | Medical waste handlers | 126 | 47.60% | 41.30% | 6.30% | 52.40% | Different healthcare workers |
| **Ethiopia** | Shiferaw (17) | BMC Res Notes. 2011 Nov 3;4:479. | DOI: 10.1186/1756-0500-4-479 | Addis Ababa | Non medical waste handlers | 126 | 31.70% | 30.90% | 0.80% | 68.30% | |
| **Gambia** | Peto (18) | BMC Infect Dis. 2014 Jan 7;14:7 | PMID: 24397793 | Multiple sites | Vaccinated young adults | 278 | 27.40% | 26.60% | 0.80% | 72.60% | Vaccination status |
| **Gambia** | Peto (18) | BMC Infect Dis. 2014 Jan 7;14:7 | PMID: 24397793 | Multiple sites | Unvaccinated young adults | 475 | 56.00% | 43.60% | 12.40% | 44.00% | |

| | | | | | | | | | | | |
|---|---|---|---|---|---|---|---|---|---|---|---|
| **Mauritania** | Mansour (19) | Journal of Medical Virology 84:1186–1198 (2012) | DOI: 10.1002/jmv.23336 | Nouakchott | Antenatal | 1020 | 66.30% | 55.60% | 10.70% | 33.70% | Different patient groups |
| **Mauritania** | Mansour (19) | Journal of Medical Virology 84:1186–1198 (2012) | DOI: 10.1002/jmv.23336 | Nouakchott | Patients | 946 | 76.50% | 58.20% | 18.30% | 23.50% | |
| **Nigeria** | Belo (20) | East Afr Med J. 2000 May;77(5):283-5. | PMID: 12858922 | Lagos | Healthcare workers (surgeons) | 167 | 61.70% | 36.00% | 25.70% | 38.30% | Different healthcare workers |
| **Nigeria** | Belo (20) | East Afr Med J. 2000 May;77(5):283-5. | PMID: 12858922 | Lagos | Admin staff (hospital) | 193 | 53.40% | 38.40% | 15.00% | 46.60% | |
| **Nigeria** | Onyekwere (21) | Niger Postgrad Med J. 2002 Sep;9(3):129-33. | PMID: 12501266 | Lagos | Diabetics | 100 | 45.00% | 25.00% | 20.00% | 55.00% | Different patient groups |
| **Nigeria** | Onyekwere (21) | Niger Postgrad Med J. 2002 Sep;9(3):129-33. | PMID: 12501266 | Lagos | Outpatients | 80 | 57.00% | 39.50% | 17.50% | 43.00% | |

| | | | | | | | | | | | |
|---|---|---|---|---|---|---|---|---|---|---|---|
| **Reunion** | Michault (22) | Bull Soc Pathol Exot. 2000 Feb;93(1):34-40. | PMID: 10774493 | Reunion | Antenatal | 1455 | 6.35% | 5.72% | 0.63% | 93.65% | Different populations |
| **Reunion** | Michault (22) | Bull Soc Pathol Exot. 2000 Feb;93(1):34-40. | PMID: 10774493 | Reunion | Prison | 100 | 10.00% | 8.00% | 2.00% | 90.00% | |
| **South Africa** | Mayaphi (23) | S Afr Med J. 2012 Feb 23;102(3 Pt 1):157-62. | PMID: 22380911 | Tshwane District Hospital, Pretoria, Gauteng province | HIV negative | 200 | 3.50% | 1.50% | 2.00% | 96.50% | HIV status |
| **South Africa** | Mayaphi (23) | S Afr Med J. 2012 Feb 23;102(3 Pt 1):157-62. | PMID: 22380911 | Tshwane District Hospital, Pretoria, Gauteng province | HIV positive | 200 | 16.00% | 9.50% | 6.50% | 84.00% | |
| **South Africa** | Mphahlele (24) | J Clin Virol. 2006 Jan;35(1):14-20. | DOI: 10.1016/j.jcv.2005.04.003 | Medunsa Campus; near Pretoria | HIV positive | 167 | 85.00% | 49.80% | 35.20% | 15.00% | HIV status |
| **South Africa** | Mphahlele (24) | J Clin Virol. 2006 Jan;35(1):14-20. | DOI: 10.1016/j.jcv.2005.04.003 | Medunsa Campus; near Pretoria | HIV negative | 128 | 82.00% | 65.80% | 16.20% | 18.00% | |
| **Togo** | Dorkenoo (25) | Med Sante Trop. 2014 Jul-Sep;24(3):266-70 | DOI: 10.1684/mst.2014.0341 | Lome | Vaccinated healthcare workers | 100 | 68.00% | 32.00% | 36.00% | 32.00% | Vaccination status |

| | | | | | | | | | | | |
|---|---|---|---|---|---|---|---|---|---|---|---|
| **Togo** | Dorkenoo (25) | Med Sante Trop. 2014 Jul-Sep;24(3):266-70 | DOI: 10.1684/mst.2014.0341 | Lome | Unvaccinated students | 50 | 88.00% | 32.00% | 56.00% | 12.00% | |
| **Uganda** | Nakwagala (26) | East Afr Med J. 2002 Feb;79(2):68-72. | PMID: 12380879 | Mulago | HIV positive | 129 | 65.10% | 47.30% | 17.80% | 34.90% | HIV status |
| **Uganda** | Nakwagala (26) | East Afr Med J. 2002 Feb;79(2):68-72. | PMID: 12380879 | Mulago | HIV negative | 129 | 41.90% | 28.80% | 13.10% | 58.10% | |
| **Uganda** | Price (27) | AIDS. 2017, March 21,Price et al, epub | DOI: 10.1097/QAD.0000000000001454 | Kampala and Entebbe | HIV positive | 2317 | 52.00% | 46.00% | 6.00% | 48.00% | Different countries |
| **Zimbabwe** | Price (27) | AIDS. 2017, March 21,Price et al, epub | DOI: 10.1097/QAD.0000000000001454 | Harare | HIV positive | 999 | 56.30% | 39.60% | 16.70% | 43.70% | |

**Supplementary Table 3: Predicted HBsAg prevalence for Northern, Eastern, Southern, Western and Central Africa with a given anti-HBc prevalence.**

Data to inform the analysis were derived from a systematic literature review (full metadata on-line (15)). Linear regression analysis data for the African regions was simulated to predict HBsAg prevalence with a given anti-HBc prevalence ranging from 5-60% and increasing in increments of 5%. Values plotted in Suppl. Fig 3.

| | **Predicted HBsAg prevalence (%) by African region** | | | | |
|---|---|---|---|---|---|
| **Anti-Hbc Prevalence (%)** | **North** | **East** | **South** | **West** | **Central** |
| **5** | 1.03 | 1.72 | 3.34 | 3.10 | 11.03 |
| **10** | 1.97 | 2.65 | 4.72 | 4.32 | 11.05 |
| **15** | 2.90 | 3.58 | 6.11 | 5.55 | 11.07 |
| **20** | 3.83 | 4.52 | 7.49 | 6.77 | 11.09 |
| **25** | 4.77 | 5.45 | 8.87 | 8.00 | 11.10 |
| **30** | 5.70 | 6.38 | 10.25 | 9.22 | 11.12 |
| **35** | 6.63 | 7.32 | 11.63 | 10.45 | 11.14 |
| **40** | 7.57 | 8.25 | 13.01 | 11.67 | 11.16 |
| **45** | 8.50 | 9.18 | 14.39 | 12.89 | 11.18 |
| **50** | 9.43 | 10.12 | 15.77 | 14.12 | 11.20 |
| **55** | 10.37 | 11.05 | 17.16 | 15.34 | 11.22 |
| **60** | 11.30 | 11.98 | 18.54 | 16.57 | 11.24 |

**Supplementary Table 4: Results of a systematic literature review to identify evidence or recommendations for use of catch up HBV vaccination in adolescents and adults in Africa.**

| Manuscript title | First author and citation | Summary comments |
|---|---|---|
| Hepatitis B virus infections in apparently healthy urban Nigerians: data from pre-vaccination tests. | Adoga (28) | Reports prevalence of HBV infection and concludes that 'The Nigerian government hepatitis B vaccination programme, which hitherto is limited to the National Childhood Immunisation Programme, should include the adult population'. |
| Decreasing the hepatitis B burden in Tunisia need more attention to adults for vaccination. | Alavian (29) | This is a letter in response to Chaouch et al (30). The authors suggest that infection is occuring in later childhood/adolescence, and advocate catch-up/adult vaccination. |
| Evidence for a change in the epidemiology of hepatitis B virus infection after nearly two decades of universal hepatitis B vaccination in South Africa. | Amponsah-Dacosta (31) | Reports wane of vaccine immunity, and the difference between HIV+ and HIV- groups. The authors raise the question 'whether the time has come to consider a pre-adolescence vaccine booster dose policy'. |
| [Study of factors influencing hepatitis B immunization coverage in 1 to 8-years-old children in the Ouidah health district in Benin in 2007]. | Bossali (32) | Study of factors influencing HBV immunization coverage in children, mothers and healthcare workers. Finding that immunization coverage decreased with age, the authors advocate 'catch-up [vaccination] sessions....in high prevalence areas' (NB. authors do not define catch up in older children vs adolescents or adults). |
| Investigating hepatitis B immunity in patients presenting to a paediatric haematology and oncology unit in South Africa. | Buchner (33) | Investigation of seroepidemiology in children in a high risk group. 40% had no immunity to HBV despite presumed vaccination. Suggests 'consider booster vaccination to the population as a whole.' |
| An update after 16 years of hepatitis B vaccination in South Africa. | Burnett (34) | Concludes on need for infant vaccine coverage, introduction of birth dose vaccine, switch to hexavalent vaccine, and consider vaccination for 12 year olds (if not vaccinated as infants). |
| Hepatitis B infection is highly endemic in Uganda: findings from a national serosurvey. | Bwogi (2) | HBsAg prevalence 10%. Risk factors: poor, un-educated, uncircumcised, ethnic group, HIV+. Conclude: 'The hepatitis B infant immunization programme should be sustained and catch-up vaccination considered for older children'. |
| Impact and long-term protection of hepatitis B vaccination: 17 years after universal hepatitis B vaccination in Tunisia. | Chaouch (30) | HBV seromarkers were checked in students. Raises the question of whether boosters in adolescence should be implemented, but doesn't currently suggest either way whether this should be undertaken. |
| Molecular epidemiology of human liver cancer: insights into etiology, pathogenesis and | Kirk (35) | Review article of HCC causes, molecular associations in the Gambia. Addresses the question of booster doses, though without reaching a firm conclusion. |

| | | |
|---|---|---|
| prevention from The Gambia, West Africa. | | |
| Observational study of vaccine efficacy 24 years after the start of hepatitis B vaccination in two Gambian villages: no need for a booster dose. | Mendy (36) | Cross sectional serological surve to determine vaccine efficacy. Efficacy against infection was 85%. Study looks at evidence for booster dose vaccine and says not helpful. |
| Long-term protection against HBV chronic carriage of Gambian adolescents vaccinated in infancy and immune response in HBV booster trial in adolescence. | van der Sande (37) | Cross-sectional study in the Gambia. Vaccine efficacy 15 years after vaccination was 67% against infection (anti-HBc positivity), and 97% against active infection (HBsAg carriage). For boosted participants anti-HBs responses were 38 IU/l prior to vaccination, 524 IU/l two weeks after boosting, and 101 IU/l after 1 year. |
| Observational study of vaccine efficacy 14 years after trial of hepatitis B vaccination in Gambian children. | Whittle (38) | Cross-sectional serological study. Vaccine-mediated antibody concentration dipped in late teen years associated with breakthrough infections. Refers to natural boosting as a result of sexual exposure in adolesence rather than booster vaccination. |
| A systematic review of hepatitis B screening economic evaluations in low- and middle-income countries. | Wright (39) | Meta-analysis of 9 studies looking at screening effectiveness in low-middle income countries. Concludes that screening with catch up vaccination for yound adults was benecifial. |

**Supplementary Fig 1: Flow chart for systematic literature review of studies reporting HBV sero-epidemiology data for countries in Africa, based on PRISMA criteria.**

The resulting metadata set is available on-line (15).

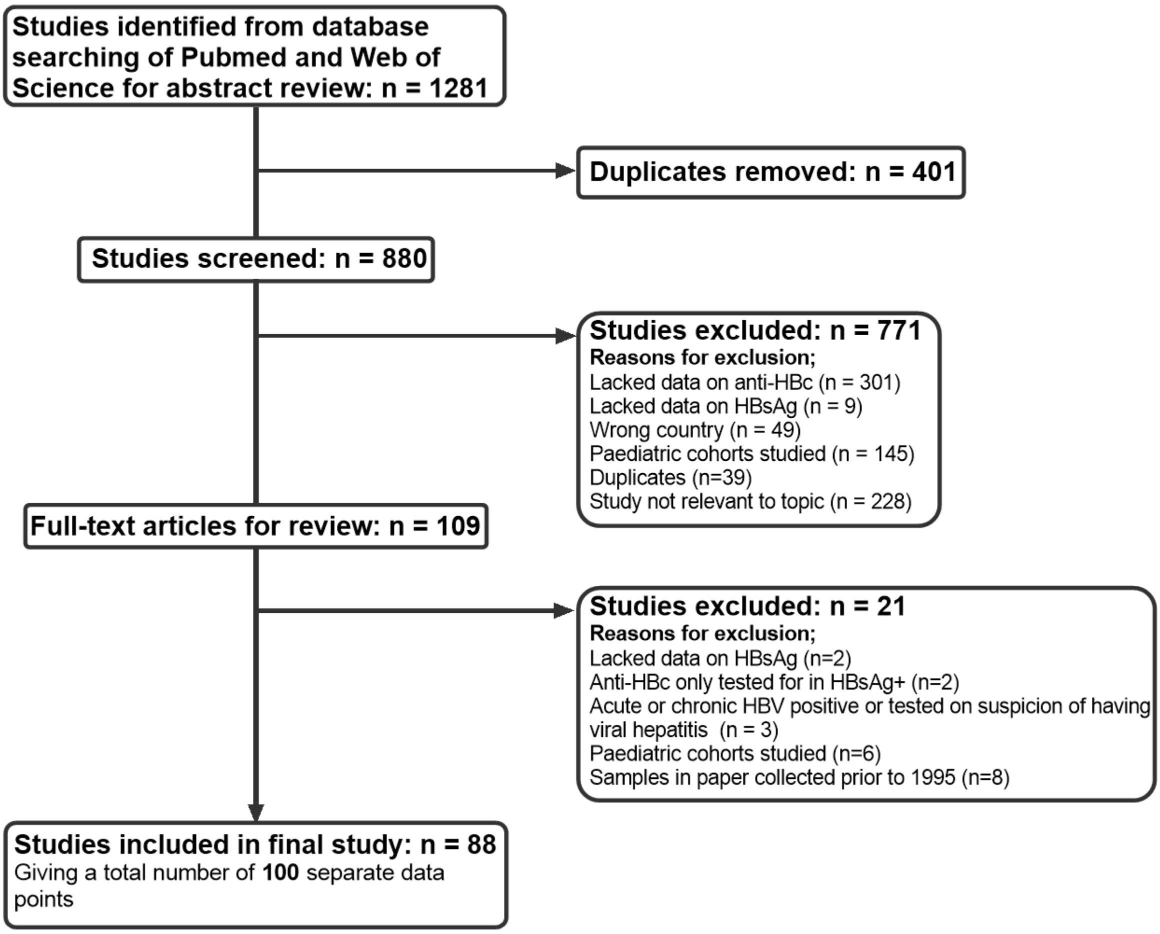

**Supplementary Fig 2: Average prevalence of anti-HBc and HBsAg in confirmed HIV-positive cohorts and all other cohorts.** Cohort characteristics were recorded for each study (Suppl data table 2). All cohorts characterised as HIV-positive (HIV+) were grouped together and compared with cohorts that were not listed at being HIV+. Box-whisker plots show the mean for each region, the interquartile ranges and the range. No significant differences were identified for either anti-HBc or HBsAg prevalence (p=0.42 and 0.16 respectively).

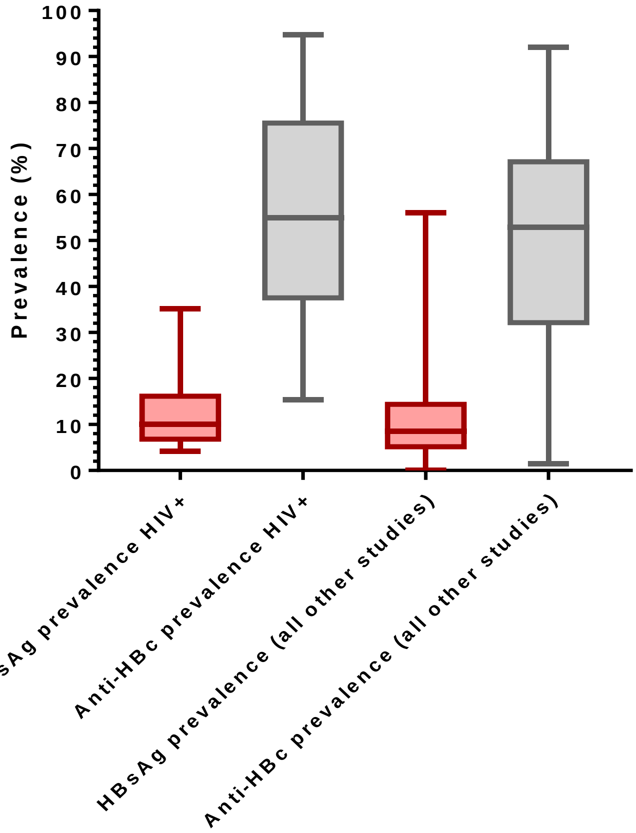

**Supplementary Fig 3: Predicted HBsAg prevalence for Northern, Eastern, Southern, Western and Central regions of Africa with a given total anti-HBc prevalence (reflecting exposure).** Linear regression analysis data for each region was simulated to predict HBsAg prevalence with a given anti-HBc prevalence ranging from 5-60% and increasing in increments of 5%. Plotted from values given in Suppl. Table 2.

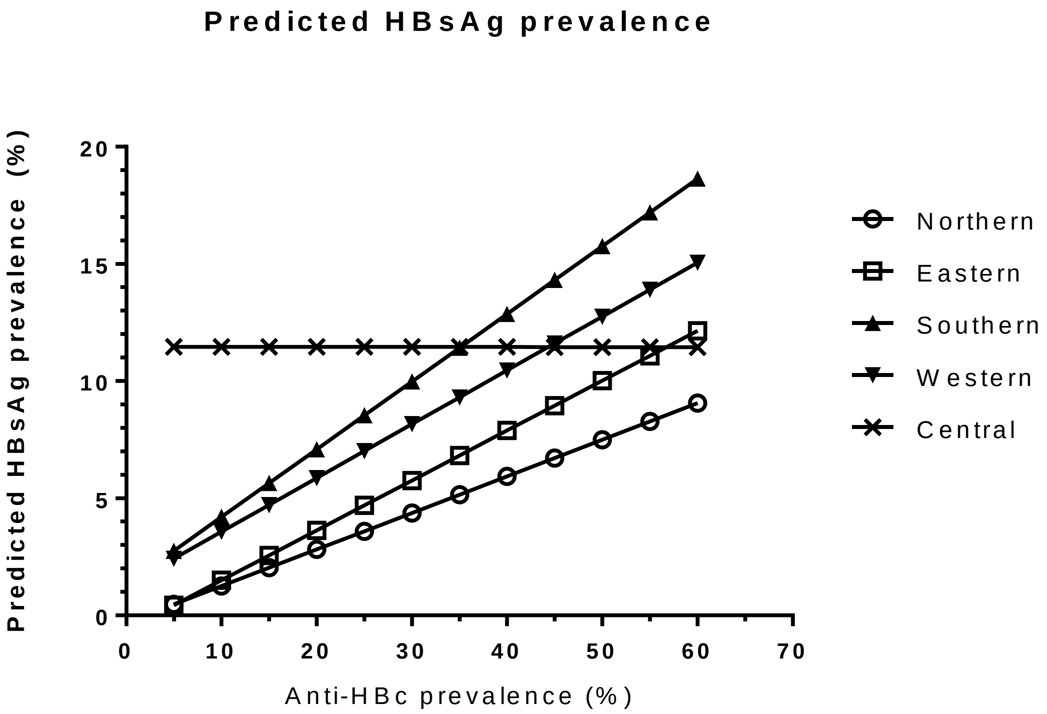
